# HASLR: Fast Hybrid Assembly of Long Reads

**DOI:** 10.1101/2020.01.27.921817

**Authors:** Ehsan Haghshenas, Hossein Asghari, Jens Stoye, Cedric Chauve, Faraz Hach

**Affiliations:** School of Computing Science, Simon Fraser University, Burnaby, Canada; Vancouver Prostate Centre, Vancouver, Canada; Faculty of Technology and Center for Biotechnology, Bielefeld University, Bielefeld, Germany; Department of Mathematics, Simon Fraser University, Burnaby, Canada; LaBRI, Université de Bordeaux, Bordeaux, France; Department of Urologic Sciences, University of British Columbia, Vancouver, Canada

**Keywords:** Hybrid assembly, Third generation sequencing, PacBio, Oxford Nanopore, Illumina

## Abstract

Third generation sequencing technologies from platforms such as Oxford Nanopore Technologies and Pacific Biosciences have paved the way for building more contiguous assemblies and complete reconstruction of genomes. The larger effective length of the reads generated with these technologies has provided a mean to overcome the challenges of short to mid-range repeats. Currently, accurate long read assemblers are computationally expensive while faster methods are not as accurate. Therefore, there is still an unmet need for tools that are both fast and accurate for reconstructing small and large genomes. Despite the recent advances in third generation sequencing, researchers tend to generate second generation reads for many of the analysis tasks. Here, we present HASLR, a hybrid assembler which uses both second and third generation sequencing reads to efficiently generate accurate genome assemblies. Our experiments show that HASLR is not only the fastest assembler but also the one with the lowest number of misassemblies on all the samples compared to other tested assemblers. Furthermore, the generated assemblies in terms of contiguity and accuracy are on par with the other tools on most of the samples.

**Availability:** HASLR is an open source tool available at https://github.com/vpc-ccg/haslr.

## 1 Introduction

Long reads (LRs) generated by third generation sequencing (TGS) technologies such as Pacific Biosciences (PacBio) and Oxford Nanopore Technologies (ONT) have revolutionized the landscape of de novo genome assembly. While LRs have higher error rate compared to short reads (SRs) generated by next generation sequencing (NGS) technologies such as Illumina, they have been shown to result in accurate assemblies given sufficient coverage. Indeed the length of TGS LRs enables the resolution of many short and mid-range repeats that are problematic when assembling genomes from SRs. Recent advances in sequencing ultra-long ONT reads have moved us closer to the complete reconstruction of entire genomes (including difficult-to-assemble regions such as centromeres and telomeres) than ever before [19]. Similarly, HiFi PacBio reads have been shown to be capable of improving the contiguity and accuracy in complex regions of the human genome [30]. These advances toward more accurate and complete genome assembly could not be achieved without the recent development of assemblers specifically tailored for LRs. These tools assemble LRs either after an error correction step [14,4] or directly without any prior error correction [17,25,12].

Although LRs are becoming more widely used for de novo genome assembly, using *hybrid* approaches (that utilize a complementary SR dataset) is still popular for several reasons: (i) SRs have higher accuracy and can be generated by Illumina sequencers at a high throughput for a lower cost; (ii) plenty of SR datasets are already publicly available for many genomes; (iii) for some basic tasks such as variant calling (SNV and short indel detection), SRs still provide better resolution due to their high accuracy which often motivates researchers to generate SRs even when LRs are in hand; and (iv) unlike PacBio assemblies whose accuracy increases with the depth of coverage thanks to their unbiased random error model [23], constructing reference quality genomes solely from ONT reads remains challenging due to biases in base calling, even with a high coverage [14,1]. As a result, hybrid assembly approaches are still useful [8,9,10].

Hybrid approaches for de novo genome assembly can be classified into three groups: (i) methods that first correct raw LRs using SRs and then build contigs using corrected LRs only (e.g. PBcR [13] and Masurca [36]); (ii) methods that first assemble raw LRs and then correct/polish the resulting draft assembly with SRs using polishing tools such as Pilon [31] and Racon [29]; and (iii) methods that first assemble SRs and then utilize LRs to generate longer contigs (e.g. hybridSPAdes [1], Unicycler [32], DBG2OLC [34], and Wengan [5]).

PBcR and Masurca correct LRs using their internal correction algorithm and then employ CABOG [21] (Celera Assembler with the Best Overlap Graph) for assembling corrected LRs. hybridSPAdes and Unicycler are similar in design. Both of these tools first use SPAdes [2] which takes SRs as input and generates an *assembly graph*, a data structure in which multiple copies of a genome segment are collapsed into a single contig (see [35] for more details). This data structure also records connections between subsequent contigs such that every region of the genome corresponds to a path in the graph. hybridSPAdes and Unicycler then align LRs to this assembly graph in order to resolve ambiguities and generate longer contigs. On the other hand, DBG2OLC first assembles contigs from SRs and maps them onto raw LRs to get a compressed representation of LRs based on SR contig identifiers, and then applies an overlap-layout-consensus (OLC) approach on these compressed LRs to assemble the genome. Since compressed LRs are much shorter compared to raw LRs, building an overlap graph from them is quicker than building it from raw LRs, due to the faster pairwise alignment. Finally, the more recent tool, Wengan, assembles short reads and then builds multiple synthetic paired-read libraries of different insert sizes from LR sequences. These synthetic paired-reads are then aligned to short read contigs and a scaffolding graph is built from the resulting alignments. In the end, the final assembly is generated by traversing proper paths of the scaffolding graph.

Among the above tools, hybridSPAdes and Unicycler have been designed specifically for bacterial and small eukaryotic genomes and do not scale for the assembly of large genomes. PBcR, Masurca, DBG2OLC and Wengan are the only hybrid assemblers that are capable of assembling large genomes, such as the human genome. However, for mammalian genomes, PBcR and Masurca require a large computational time and cannot be used without a computing cluster. DBG2OLC is faster due to its use of compressed LRs. Wengan is also a fast assembler and can be used for assembling large genomes in a reasonable time.

In this paper, we introduce HASLR, a fast hybrid assembler that is capable of assembling large genomes. HASLR, similar to hybridSPAdes, Unicycler, and Wengan builds SR contigs using a fast SR assembler (i.e. Minia). Then it builds a novel data structure called *backbone graph* to put short read contigs in the order expected to appear in the genome and to fill the gaps between them using consensus of long reads. Based on our results, HASLR is the fastest between all the assemblers we tested, while generating the lowest number of mis-assemblies. Furthermore, it generates assemblies that are comparable to the best performing tools in terms of contiguity and accuracy. HASLR is also capable of assembling large genomes using less time and memory than other tools.

## 2 Methods

The input to HASLR is a set of long reads (LRs) and a set of short reads (SRs) from the same sample, together with an estimation of the genome size. HASLR performs the assembly using a novel approach that rapidly assembles the genome without performing all-vs-all LR alignments. The core of HASLR is to first assemble contigs from SRs using an efficient SR assembler and then to use LRs to find sequences of such contigs that represent the backbone of the sequenced genome.

### 2.1 Obtaining unique short read contigs

HASLR starts by assembling SRs into a set of *short read contigs* (SRCs), denoted by *C*. Assembly of SRs is a well-studied topic and many efficient tools have been specifically designed for that purpose. These tools use either a de Bruijn graph [28,3] or an OLC strategy (based on an overlap graph or a string graph) [27,22] to assemble the genome by finding “proper” paths in these graphs.

Next, HASLR identifies a set *U* of unique contigs (UCs), those SRCs that are likely to appear in the genome only once. In order to do this, for every SRC, *c_i_*, the mean *k*-mer frequency, *f* (*c_i_*), is computed as the average *k*-mer count of all *k*-mers present in *c_i_*. Note that the value of *f* (*c_i_*) is proportional to the depth of coverage of *c_i_*. Assuming longer contigs are more likely to come from unique regions, their mean *k*-mer frequency can be a good indicator for identifying UCs. Let *LC_q_* ⊆ *C* be the set of *q* longest SRCs in *C*, and *f_avg_*, *f_std_* be the average and standard deviation of {*f* (*c*) | *c* ∈ *LC_q_*}. Then, the set of unique contigs is defined as *U* = {*u* | *u* ∈ *C* and *f* (*u*) ≤ *f_avg_* + 3*f_std_*}. Our empirical results show that this approach can identify UCs with high precision and recall (see Supplementary Section S4 for more details).

### 2.2 Construction of backbone graph

The backbone graph encodes potential adjacencies between unique contigs and thus presents a large-scale map of the genome, albeit, with some level of ambiguity. Using the backbone graph, HASLR finds paths of unique contigs representing their relative order and orientation in the sequenced genome. These paths are later transformed into the assembly.

Formally, given a set of UCs, *U* = {*u*_1_, *u*_2_, …, *u_|U|_*}, and a set of LRs, *L* = {*l*_1_, *l*_2_, …, *l_|L|_*}, HASLR builds the backbone graph *BBG* as follows. First, UCs are aligned against LRs. Each alignment can be encoded by a 7-tuple (*rbeg, rend, uid, ustrand, ubeg, uend, nmatch*) whose elements respectively denote the start and end positions of the alignment on the LR, the index of the UC in *U*, the strand of the alignment (+ or −), the start and end position of the alignment on the UC, and the number of matched bases in the alignment. Let 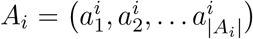 be the list of alignments of UCs to *l_i_*, sorted by *rend*.

Note that alignments in *A_i_* may overlap due to relaxed alignment parameters in order to account for the high sequencing error rate of LRs. Thus, in the next step we aim to select a subset of non-overlapping alignments whose total identity score – defined as the sum of the number of matched bases – is maximal. Let *S_i_*(*j*) be the best subset among the first *j* alignments, i.e. the non-overlapping subset of these *j* alignments with maximal total identity score. *S_i_*(*j*) can be calculated using the following dynamic programming formulation:

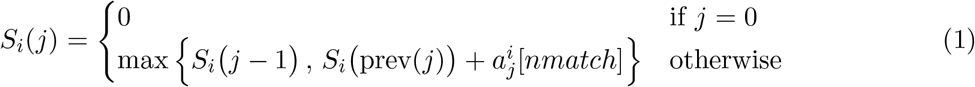

where prev(*j*) is the largest index *z < j* such that 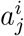 and 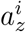 are non-overlapping alignments. By calculating *S_i_*(|*A_i_*|) and backtracking, we obtain a sorted sub-list 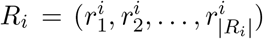 of non-overlapping alignments with maximal total identity score, which we call the *compact representation* of read *l_i_*. Note that since the input list is sorted, prev(.) can be calculated in logarithmic time which makes the time complexity of this dynamic programming *O*(|*A_i_*| log |*A_i_*|).

The backbone graph is a *directed* graph *BBG* = (*V, E*). The set of nodes is defined as 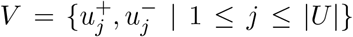 where 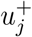 and 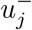 represent the forward and reverse strand of the UC *u_j_*, respectively. The set of edges is defined as the oriented adjacencies between UCs implied by the compact representations of LRs. Formally each edge is represented by a triplet (*u_h_*, *u_t_*, *supp*) where *u_h_*, *u_t_* ∈ *V* and *supp* is the set of indices of LRs supporting the adjacency between *u_h_* and *u_t_*; these triplets are obtained as follows:

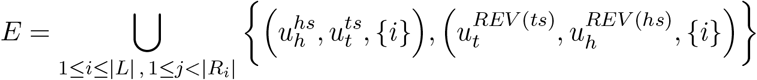

where 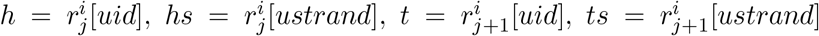, *REV* (+) = −, and *REV* (−) = +. Figure 1 illustrates the construction of the backbone graph edges for several combinations of UC alignments on LRs.

**Fig. 1:**
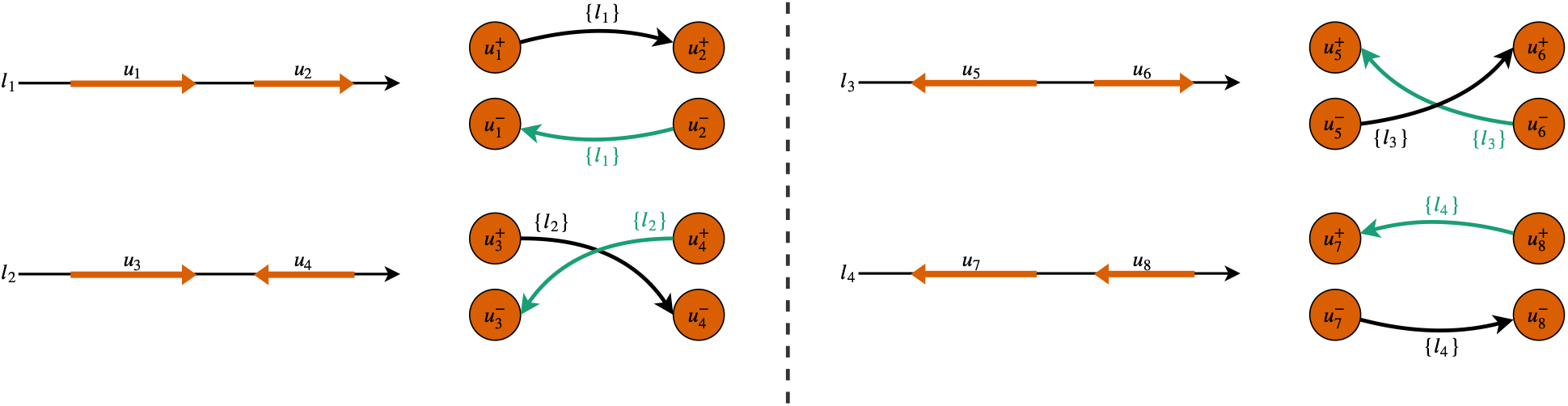
Possible orientations of aligning two unique contigs to a long read. The direction of contigs aligned to long reads shows the strand of their corresponding sequence. These directions guide us to find the proper edge type. The set of long reads supporting each edge is shown as its label.

At the end of this stage, the resulting backbone graph is a multi-graph as there can be multiple edges between two nodes with different *supp*. In order to make it easier to process the backbone graph, we convert it into a simple graph by merging *supp* of all edges between every pair of nodes into a set of supporting LRs.

### 2.3 Graph cleaning and simplification

Ideally, with accurate identification of UCs and correct alignment of UCs onto LRs, the backbone graph for a *haploid genome* will consist of a set of connected components, each of which is a *simple path* of nodes. In practice, this ideal case does not happen – mainly due to sequencing errors, wrong UC to LR alignments, and chimeric reads. As a result, some artifactual branches might exist in the backbone graph forming structures known as *tips* and *bubbles*. Tips are dead-end simple paths with a small length. Bubbles are formed when two disjoint simple paths with similar length occur between two nodes.

We clean the backbone graph *BBG* in two stages. First, in order to reduce the effect of wrong UC to LR alignments, we remove all edges *e* such that |*e*[*supp*]| < *minSupp*, for a given parameter *minSupp*. Second, the graph is simplified by using tip and bubble removal algorithms. There exist well-known algorithms for these tasks that are commonly used in assemblers [35,2,22]. Note that our tip and bubble removal procedures require an estimation of the length of simple paths. Such estimation can be obtained from the length of UCs corresponding to the nodes contained in a simple path as well as the average length of all LR subsequences that are supporting edges between consecutive nodes (see Supplementary Section S5 for more details). We denote by *G* the cleaned and simplified backbone graph.

### 2.4 Generating the assembly

The principle behind the construction of the assembly is that each simple path in the cleaned backbone graph *G* is used to define a contig of this assembly. Suppose *P* = (*v*_1_, *e*_12_, *v*_2_, *e*_23_, *v*_3_, …, *v_n_*) is a simple path of *G*. Although we already have the DNA sequence for each UC corresponding to each node *v_i_*, the DNA sequence of the resulting contig cannot be obtained immediately. This is due to the fact that at this stage the subsequence between *v_i_* and *v_i_*_+1_ is unknown for each 1 ≤ *i < n*. Here, we explain how these missing subsequences are reconstructed.

For simplicity, suppose we would like to obtain the subsequence between the pair *v*_1_ and *v*_2_ in *P*. Note that by construction, *e*_12_[*supp*] contains all LRs supporting *e*_12_. We can extract a compact representation of all those LRs and align them to *P* using a Longest Common Subsequence (LCS) dynamic programming algorithm forbidding mismatches (only gaps are allowed). We implemented this LCS algorithm in a way that takes into account the strand of UCs in *P* (recall that 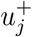 and 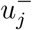 correspond to the forward and reverse strand of *u_j_* respectively). At this point, we can extract the subsequence between *v*_1_ and *v*_2_ from each LR in *e*_12_[*supp*]. To do this, we find the region of UCs corresponding to *v*_1_ and *v*_2_ that are aligned to all LRs in *e*_12_[*supp*]. Using the alignment transcript (i.e. CIGAR string) the unaligned coordinate of each long read is calculated (see Figure 2 for a toy example). By computing the consensus sequence of the extracted subsequences, we obtain *cns*_12_. Therefore, the DNA sequence corresponding to *P* can be obtained via *CONCAT* (*u*_1_, *cns*_12_, *u*_2_, *cns*_23_, *u*_3_, …, *u_n_*) where *CONCAT* (.) returns the concatenated DNA sequence of all its arguments.

**Fig. 2:**
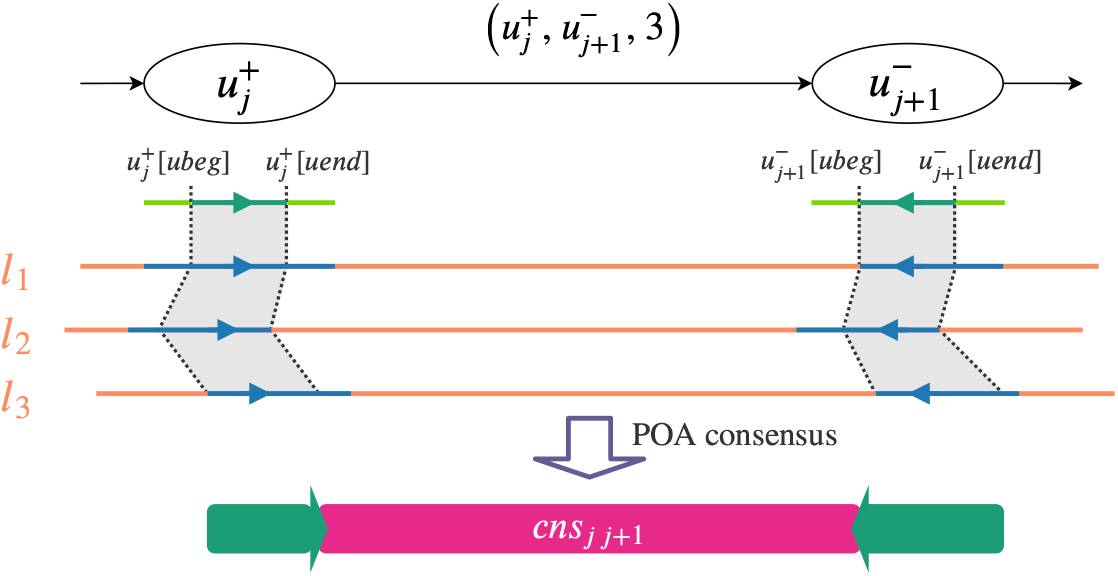
Example of an edge in backbone graph and its corresponding long read alignments. Partial Order Alignment (POA) is used in constructing the consensus sequence (see subsection 2.5)

In order to generate the assembly, HASLR extracts all the simple paths in the cleaned backbone graph *G* and constructs the corresponding contig for each of them as explained above. It is important to note that each simple path *P* has a *twin* path *P′* which corresponds to the reverse complement of the contig generated from *P*. Therefore, during our simple path extraction procedure, we ensure to not use twin paths to avoid redundancy.

### 2.5 Methodological remarks

#### Rationale for using unique short read contigs

Here, we clarify the motivation for choosing only unique SRCs as the nodes of the backbone graph. Repetitive genomic regions cause complexities in assembly graphs. The same complexity is reflected in our backbone graph. Repetitive SRCs would cause branching in the backbone graph and in fact, building the backbone graph using all SRCs could result in a very tangled graph. Figure 3 illustrates the difference between a backbone graph built on all SRCs with one built only on unique SRCs on a yeast genome. As it can be seen, using only unique SRCs for building the backbone graph resolves many of the complexities and ambiguities in the graph. However, it is important to note that excluding non-unique SRCs could potentially result in a more fragmented graph (some chromosomes are split into multiple paths rather than a single one) and assembly.

**Fig. 3:**
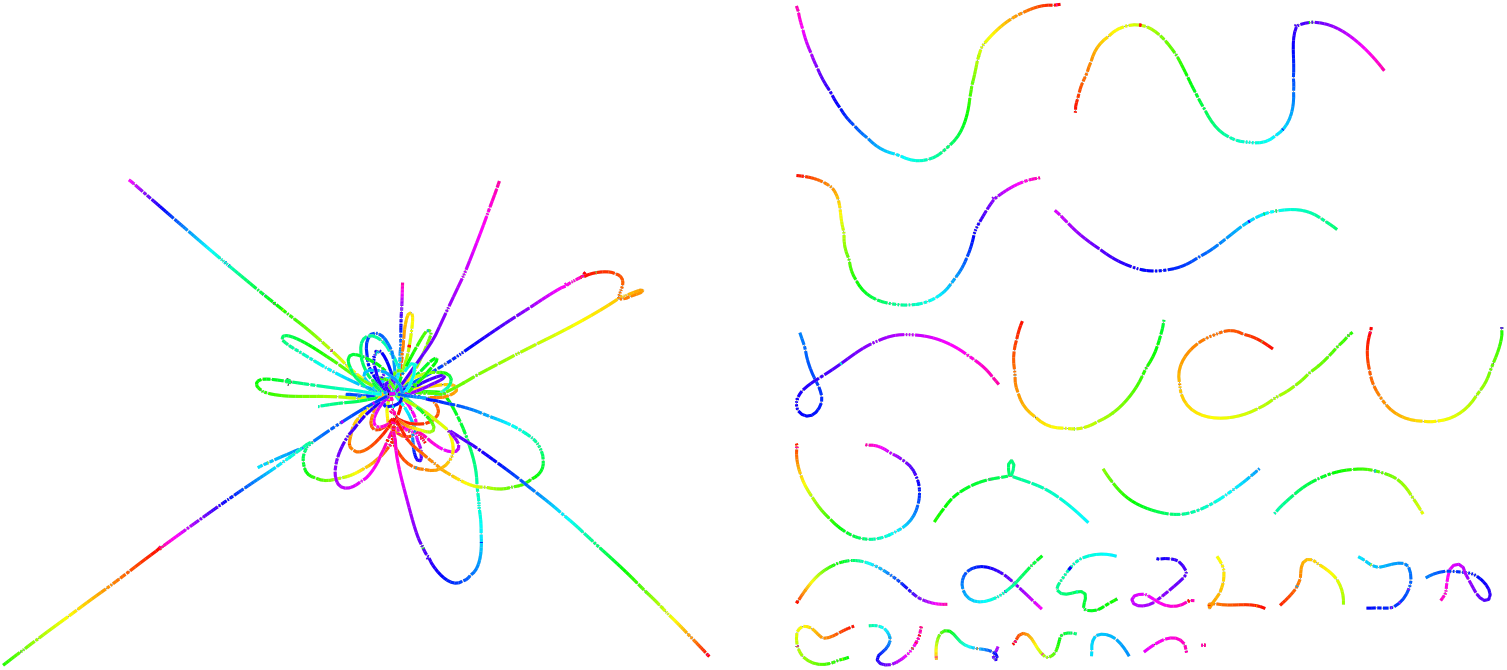
Two backbone graphs built from a real PacBio dataset sequenced from a yeast genome. Each graph is visualized with Bandage [33] and colored using its rainbow coloring feature. Each chromosome is colored with a full rainbow spectrum. (Left) Tangled graph built from all SRCs. (Right) Untangled graph built from unique SRCs.

#### Backbone graph vs. assembly graph

It is important to note that the backbone graph is not an assembly graph per se, for two reasons. First, the regions between each pair of connected unique SRCs are not present in the graph. These missing regions are obtained by calculating the consensus of LR subsequences between each pair of unique SRCs. Second, unlike assembly graphs, there are some segments of the genome that cannot be translated to a path in the backbone graph. This is due to the potential fragmentation that was mentioned earlier.

#### Implementation details

(i) HASLR utilizes a SR assembler to build its initial SRCs. However, a higher quality assembly that has fewer misassemblies is preferred. For this purpose, HASLR utilizes Minia [3] to assemble SRs into SRCs. Based on our experiments, Minia can generate a high quality assembly quickly using a small memory footprint. (ii) For finding UCs, HASLR calculates mean *k*-mer frequencies with a small value of *k* (default *k* = 55). This information can be easily obtained by performing a *k*-mer counting on the SR dataset (for example using KMC [11]) and calculating the average *k*-mer count of all *k*-mers present in each SRC. Nevertheless, usually assemblers automatically provide such information (e.g Minia and SPAdes). HASLR takes *k*-mer frequencies reported by Minia for this task. (iii) HASLR uses only longest 25× coverage of long reads for building the backbone graph which are extracted based on the given expected genome size. (iv) In order to align UCs to LRs, HASLR employs minimap2 [18]. (v) Graph cleaning is done with *minSupp* = 3 meaning that any edge that is supported with less than 3 LRs is discarded. (vi) Finally, consensus sequences are obtained using the Partial Order Alignment [16,15] (POA) algorithm implemented in the SPOA package [29]. We have provided the versions of the tools and the parameters that are used to execute them in Supplements S2 and S3, respectively.

## 3 Results

We evaluated the performance of HASLR on both simulated and real datasets. We selected five hybrid assemblers (hybridSPAdes [1], Unicycler [32], DBG2OLC [34], Masurca [36] and Wengan [5]) as well as two non-hybrid methods (Canu [14] and wtdbg2 [25]). All experiments were performed on isolated nodes of a cluster (i.e. no other simultaneous jobs were allowed on each node). Each node runs CentOS 7 and is equipped with 32 cores (2 threads per core; total of 64 CPUs) Intel(R) Xeon(R) processors (Gold 6130 @ 2.10GHz) and 720 GB of memory. Each tool was run with their recommended settings. See supplementary Sections S2 and S3 for more details about the versions of tools and the employed commands. Note that for wtdbg2, we used the provided `wtdbg2.pl` wrapper which automatically performs a polishing step using the embedded polishing module.

For each experiment, assemblies were evaluated by comparing against their corresponding reference genome using QUAST [20]. QUAST reports on a wide range of assembly statistics but we are mostly interested in misassemblies, NGA50 and rate of small errors (mismatch or indel). QUAST detects and reports misassemblies when a contig cannot align to the reference genome as a single continuous piece. Misassemblies indicate structural assembly errors. For computing NGA50, unlike N50 and NG50, only segments of assembled contigs that are aligned to the reference genome are considered. In addition, QUAST breaks contigs with extensive misassemblies before calculation of NGA50. Therefore, NGA50 is a good indicator of the contiguity of the assembly, while taking misassemblies into consideration.

### 3.1 Experiment on simulated dataset

We evaluated all the selected methods on 4 simulated datasets, namely *E. coli*, yeast, *C. elegans* and human, to provide a wide range of genome sizes and complexity. For each genome, we used ART [7] to simulate 50× coverage short Illumina reads (2×150 bp long, 500 bp insert size mean, and 50 bp insert size deviation) using the Illumina HiSeq 2000 error model. We also simulated 50× coverage long PacBio reads using PBSIM [24]. In order to capture the characteristics of real datasets, a set of PacBio reads generated from a human genome (See Supplementary Section S1.1 for details) with P6-C4 chemistry was passed to PBSIM via option `--sample-fastq`. This enables PBSIM to sample the read length and error model from the real long reads.

Table 1, shows the QUAST metrics calculated for assemblies generated by different tools. As it can be seen, HASLR generates assemblies with the lowest number of misassemblies in all datasets. It is important to note that since reads are simulated from the same reference used for this assessment, any misassembly reported by QUAST is indeed a structural assembly mistake. In terms of the contiguity, HASLR achieves NGA50 on par with other tools for all datasets except for *C. elegans* where Canu shows an NGA50 twice larger than others tools. On the human dataset, HASLR generates the most contiguous assembly with an NGA50 of 17.03 Mb and only 2 extensive misassemblies, although at the price of a lower genome fraction (see Discussion). In addition, HASLR is the fastest assembler across the board. wtdbg2 has a comparable speed but generates lower quality assemblies, both in terms of misassemblies and mismatch/indel rate.

**Table 1:**
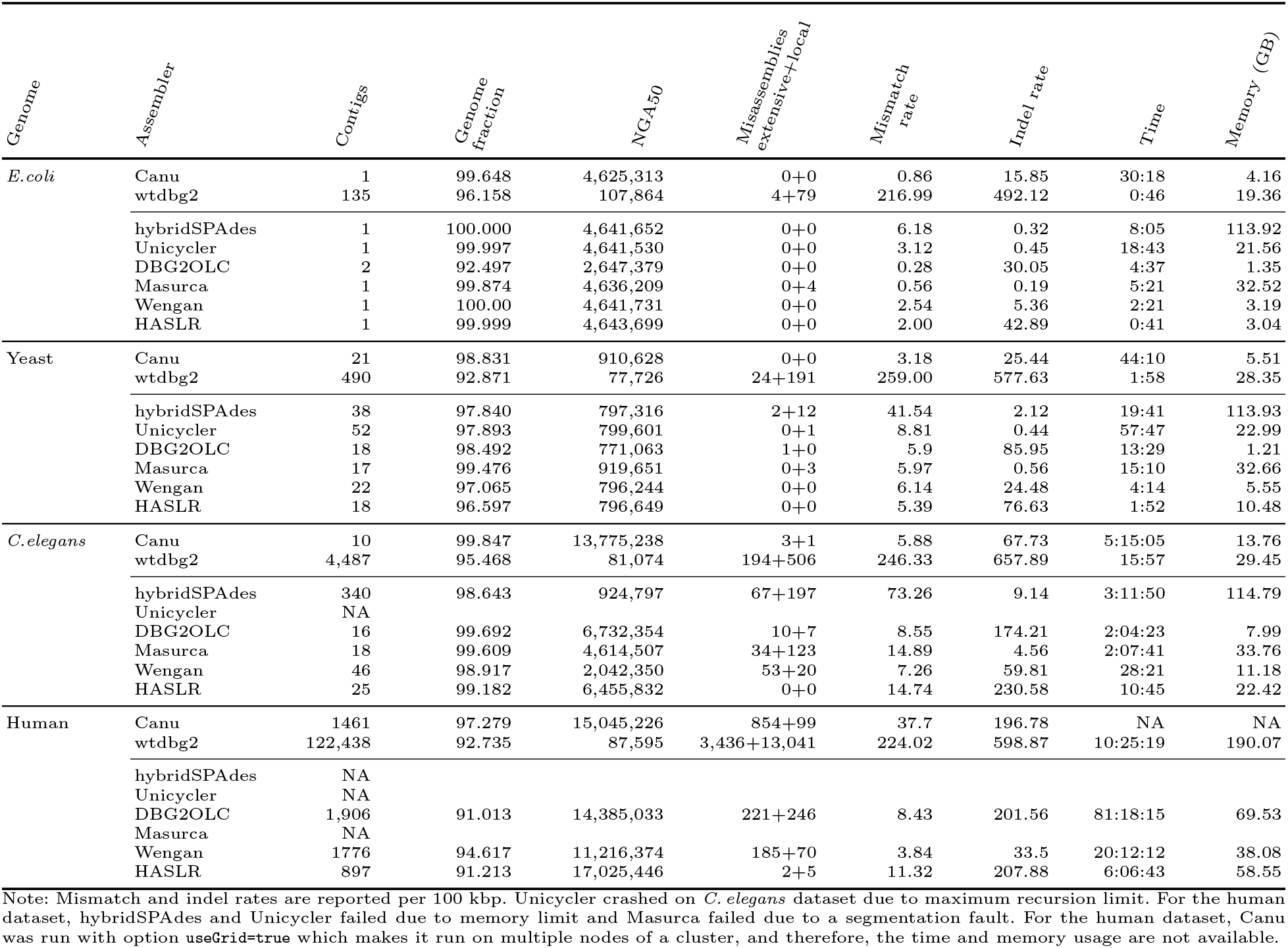
Comparison between draft assemblies obtained by different tools on simulated data.

It is particularly interesting to compare HASLR with hybridSPAdes, Unicycler and Wengan, since they share similar design in that they connect short read contigs rather than explicitly assembling long reads. In addition, Wengan uses short read contigs generated by Minia, similar to HASLR. hybridSPAdes and Unicycler do not scale for large genomes as they have been designed for small and bacterial genomes. On *C. elegans* dataset, HASLR gives significantly more contiguous assembly than hybridSPAdes and Wengan without any structural assembly error. For the human dataset, HASLR has a higher NGA50 while generating significantly less misassemblies.

Note that, HASLR does not employ any polishing step neither internally nor externally. Thus, the indel rate of the draft assemblies generated by HASLR is less than desirable. However, these types of local assembly erros can be easily addressed through a polishing step as shown in Table S3. With a single round of polishing, both indel and mismatches rates match the other tools in two datasets.

### 3.2 Experiment on real dataset

To compare the performance of HASLR on real dataset with other tools, we tested them on 4 publicly available datasets, *E. coli*, yeast, *C.elegans*, and human. Table 2 contains details about these real datasets (see Table S1 for the availability of each dataset). Similar to simulated datasets, on real dataset HASLR generates less misassembly compared to other assemblers while remaining the fastest. Compared to other hybrid assemblers, HASLR performs similar or better in terms of contiguity, while stands behind self-assembly tools with a lower NGA50.

**Table 2:** Statistics of real long read datasets

| Dataset | Technology | N50 length | Estimated coverage | Total size (Gb) | Aligned size (Gb) | Avg. alignment identity (%) |
| --- | --- | --- | --- | --- | --- | --- |
| *E. coli*<br>(K-12 MG1655) | ONT R9.4 | 63,747 | 1,080 | 5.01 | 4.31 | 85.03 |
|  | Illumina | 2×151 | 372 | 1.73 | - | - |
| Yeast<br>(S288C) | PacBio | 8,561 | 132 | 1.61 | 1.42 | 86.90 |
|  | Illumina | 2×150 | 82 | 1.00 | - | - |
| *C. elegans*<br>(Bristol) | PacBio | 16,675 | 47 | 4.73 | 4.32 | 87.43 |
|  | Illumina | 2×100 | 67 | 6.76 | - | - |
| Human<br>(CHM1) | PacBio | 19,960 | 59 | 182.51 | 163.51 | 85.85 |
|  | Illumina | 2×151 | 41 | 127.76 | - | - |

Note: Alignment statistics were obtained by aligning long reads against their reference genome using `lordFAST` [6].

**Table 3:** Comparison between assemblies obtained by different tools on real data

| *Dataset* | *Assembler* | *Contigs* | *Genome fraction* | *NGA50* | *Misassemblies extensive+local* | *Mismatch rate* | *Indel rate* | *Time* | *Memory (GB)* |
| --- | --- | --- | --- | --- | --- | --- | --- | --- | --- |
| *E. coli* (ONT) | Canu | 1 | 99.976 | 3,647,271 | 2+6 | 108.85 | 1254.40 | 702:57:07 | 32.39 |
|  | wtdbg2 | 9 | 79.114 | 141,474 | 38+72 | 245.82 | 1501.74 | 4:57 | 28.05 |
|  | hybridSPAdes | 15 | 99.964 | 3,863,268 | 2+7 | 7.16 | 0.50 | 3:38:13 | 114.29 |
|  | Unicycler | NA |  |  |  |  |  |  |  |
|  | DBG2OLC | 1 | 99.950 | 3,539,045 | 3+4 | 46.86 | 335.82 | 8:25 | 8.74 |
|  | Masurca | 1 | 99.988 | 3,892,134 | 3+7 | 2.82 | 0.50 | 30:28 | 32.66 |
|  | Wengan | 3 | 99.998 | 3,346,596 | 3+2 | 4.74 | 9.24 | 20:02 | 14.37 |
|  | HASLR | 2 | 99.992 | 3,970,011 | 2+2 | 22.62 | 79.85 | 3:18 | 5.78 |
| Yeast (PacBio) | Canu | 23 | 99.724 | 739,932 | 29+2 | 8.85 | 7.99 | 1:00:19 | 5.97 |
|  | wtdbg2 | 28 | 97.668 | 640,895 | 20+3 | 10.65 | 27.17 | 3:04 | 16.26 |
|  | hybridSPAdes | 61 | 97.207 | 436,584 | 28+20 | 44.77 | 3.71 | 20:58 | 114.09 |
|  | Unicycler | 51 | 97.555 | 531,185 | 15+5 | 15.13 | 4.22 | 2:09:27 | 36.90 |
|  | DBG2OLC | 24 | 63.275 | 229,397 | 25+10 | 28.37 | 58.43 | 9:51 | 0.99 |
|  | Masurca | 24 | 99.262 | 538,374 | 30+8 | 11.83 | 5.85 | 23:15 | 32.69 |
|  | Wengan | 29 | 96.258 | 528,763 | 14+10 | 11.86 | 34.29 | 6:38 | 8.64 |
|  | HASLR | 28 | 95.735 | 530,856 | 11+5 | 8.13 | 100.64 | 2:25 | 11.30 |
| *C. elegans* (PacBio) | Canu | 172 | 99.665 | 561,201 | 723+596 | 65.28 | 58.82 | 4:15:23 | 11.62 |
|  | wtdbg2 | 288 | 98.994 | 561,292 | 329+596 | 26.82 | 79.72 | 14:13 | 21.19 |
|  | hybridSPAdes | 2,336 | 96.72 | 84,003 | 633+638 | 108.04 | 15.96 | 2:47:32 | 74.11 |
|  | Unicycler | 858 | 97.102 | 139,992 | 940+692 | 58.36 | 45.47 | 23:49:29 | 105.06 |
|  | DBG2OLC | 206 | 99.100 | 421,196 | 546+383 | 44.75 | 80.61 | 2:34:44 | 11.36 |
|  | Masurca | 216 | 97.013 | 471,366 | 368+504 | 49.20 | 23.50 | 1:57:49 | 33.48 |
|  | Wengan | 270 | 93.341 | 341,861 | 308+336 | 35.75 | 121.11 | 45:45 | 33.48 |
|  | HASLR | 261 | 97.431 | 453,631 | 259+331 | 26.08 | 140.40 | 15:35 | 17.93 |
| CHM1 (PacBio) | Canu | 2,110 | 96.084 | 2,329,909 | 6,715+7,048 | 145.81 | 120.69 | 689:26:01 | 70.44 |
|  | wtdbg2 | 3,723 | 92.896 | 2,081,842 | 3,535+6,286 | 118.45 | 72.54 | 11:35:22 | 202.41 |
|  | hybridSPAdes | NA |  |  |  |  |  |  |  |
|  | Unicycler | NA |  |  |  |  |  |  |  |
|  | DBG2OLC | 2,118 | 95.547 | 1,599,466 | 3,718+8,690 | 116.81 | 116.89 | 78:21:08 | 64.94 |
|  | Masurca | 3,781 | 93.782 | 1,761,291 | 4,984+7,491 | 180.83 | 57.53 | 350:35:59 | 225.63 |
|  | Wengan | 4,474 | 88.948 | 875,489 | 2,771+7,577 | 115.65 | 160.71 | 18:19:47 | 112.73 |
|  | HASLR | 1,469 | 92.664 | 1,699,092 | 2,097+7,661 | 113.06 | 281.74 | 6:32:33 | 60.75 |

Note: Mismatch and indel rates are reported per 100 kbp. hybridSPAdes and Unicycler failed on human genome datasets due to memory limit. Unicycler did not finish on *E. coli* dataset within two weeks.

For real datasets, we further evaluated the accuracy of assemblies by performing gene completeness analysis using BUSCO [26], which quantifies gene completeness using single-copy orthologs. Table 4 shows the results of BUSCO on *E. coli*, yeast, and *C. elegans*. We were unable to obtain BUSCO results for the human genome due to a high run time requirement.

**Table 4:** Gene completeness analysis

| Dataset | Assembler | Complete (%) | Complete single copy (%) | Complete duplicate (%) | Fragmented (%) | Missing (%) | Total BUSCO groups |
| --- | --- | --- | --- | --- | --- | --- | --- |
| *E. coli* (ONT) | Canu | 4.1 | 4.1 | 0.0 | 16.8 | 79.1 | 440 |
|  | wtdbg2 | 1.8 | 1.8 | 0.0 | 9.1 | 89.1 | 440 |
|  | hybridSPAdes | 100.0 | 99.5 | 0.5 | 0.0 | 0.0 | 440 |
|  | Unicycler | NA |  |  |  |  |  |
|  | DBG2OLC | 35.9 | 35.7 | 0.2 | 33.0 | 31.1 | 440 |
|  | Masurca | 99.7 | 98.6 | 1.1 | 0.0 | 0.3 | 440 |
|  | Wengan | 100.0 | 99.5 | 0.5 | 0.0 | 0.0 | 440 |
|  | HASLR | 97.8 | 97.3 | 0.5 | 1.6 | 0.6 | 440 |
| Yeast (PacBio) | Canu | 96.6 | 94.8 | 1.8 | 0.2 | 3.2 | 2137 |
|  | wtdbg2 | 88.4 | 86.8 | 1.6 | 0.8 | 10.8 | 2137 |
|  | hybridSPAdes | 96.6 | 94.8 | 1.8 | 0.1 | 3.3 | 2137 |
|  | Unicycler | 96.4 | 94.7 | 1.7 | 0.1 | 3.5 | 2137 |
|  | DBG2OLC | 57.1 | 56.5 | 0.6 | 0.5 | 42.4 | 2137 |
|  | Masurca | 96.3 | 94.1 | 2.2 | 0.1 | 3.6 | 2137 |
|  | Wengan | 96.5 | 94.9 | 1.6 | 0.0 | 3.5 | 2137 |
|  | HASLR | 95.8 | 94.4 | 1.4 | 0.1 | 4.1 | 2137 |
| *C. elegans* (PacBio) | Canu | 97.4 | 96.8 | 0.6 | 1.1 | 1.5 | 3131 |
|  | wtdbg2 | 97.1 | 96.5 | 0.6 | 1.3 | 1.6 | 3131 |
|  | hybridSPAdes | 96.4 | 95.8 | 0.6 | 1.3 | 2.3 | 3131 |
|  | Unicycler | 97.7 | 97.1 | 0.6 | 0.7 | 1.6 | 3131 |
|  | DBG2OLC | 97.5 | 95.8 | 1.7 | 0.6 | 1.9 | 3131 |
|  | Masurca | 95.5 | 94.1 | 1.4 | 0.4 | 4.1 | 3131 |
|  | Wengan | 91.6 | 91.1 | 0.5 | 0.9 | 7.5 | 3131 |
|  | HASLR | 97.1 | 96.7 | 0.4 | 0.8 | 2.1 | 3131 |

Note: We used enterobacterales\_odb10, saccharomycetes\_odb10, and nematoda\_odb10 gene sets for assessing gene completeness of *E. coli*, Yeast, and *C. elegans* assemblies, respectively. We were not able to obtain the gene completeness results for the human dataset due to time restrictions.

Another observation is that for some experiments, HASLR does not perform as well as others in terms of genome fraction (see Discussions for more details). However, our gene completeness analysis shows that HASLR is on par with other tools based on BUSCO gene completeness measure (see Table 4). Note that very low gene completeness of Canu, wtdbg2, and DBG2OLC on *E. coli* dataset could be due to high indel rates of their assemblies.

## 4 Discussion

HASLR introduces the notion of backbone graph for hybrid genome assembly. This enables HASLR to keep up with increasing throughput of LR sequencing technologies while remaining time and memory efficient. The high speed of HASLR is due to two reasons; (i) HASLR uses the fast SPOA consensus module rather than normal POA implementation, and (ii) HASLR uses only longest 25× coverage of LRs for assembly. Assemblies generated by HASLR are similar to those generated by best-performing tools in terms of contiguity while having the lowest number of misassemblies. In other words, we prefer to remain conservative in resolving ambiguous regions without strong signal rather than aggressively resolving them to generate longer contigs and possibly generating misassemblies. However the conservative nature of HASLR does not implies it compromises on assembling complex regions. Every complex region that is covered by a sufficient number of LRs, together with its flanking unique SR contigs, would be resolved. In fact, based on our manual inspections, there are regions that HASLR assembles properly but all other tools either misassemble or generate fragmented assembly (see supplementary Section S7 for visual examples of such cases).

There are a number of future directions that are planned for future releases of HASLR. First, compared to other tools, HASLR usually has a higher indel rate. Note that most of the small local assembly mistakes (including mismatch and indel errors) can be fixed by further polishing. But since a large portion of the assembled genome is built from SRCs, a polishing module could be specifically designed for HASLR that only polishes the regions between unique SRCs which have been generated using SPOA. This would enable a faster polishing phase.

HASLR sometimes generates assemblies with relatively lower genome fraction than other tools. This is more clear when we compare it against Canu, especially on a large and complex genome like the human genome. The main reason is the lack of unique SRCs in a large region. This limitation could be mitigated by extracting unused LRs and assembling them in an OLC fashion (e.g. using miniasm [17]). Note that only a small portion of LRs are unused compared to the original input dataset. As a result, using an OLC approach for such a small set of LRs should not affect the total running time significantly.

One of the main bottlenecks of OLC-based assembly approach in terms of speed is that they require to find all overlaps between input reads. Recent LR assemblers have tried to speed up this process by using minimizers [17,14] or compressed representation of LRs [25] techniques. However, an all-versus-all alignment is still required in order to generate such a graph. In fact, OLC-based assemblers can use HASLR (or the idea of backbone graph assembly) as a first step before performing the computationally expensive all-vs-all alignment step.

An important factor in the contiguity of assemblies generated by HASLR is the length of reads. Obviously, longer reads would generate a more connected and resolved backbone graph. With the recent advancements in the Nanopore technology and the introduction of ultra-long Nanopore reads (whose length can go beyond 1 Mbp), one can expect to get much more contiguous assemblies. Therefore, supporting ultra-long ONT reads is an important feature to address in the future.

Finally, heterozygosity-aware consensus calling of subreads falling between to unique SRCs is one of our main future directions. This would be possible via clustering of subreads that fall between consecutive unique SRCs into two groups and performing consensus calling for each group separately. This would enable HASLR to perform phased assembly of diploid genomes.

## References

1. Antipov, D., Korobeynikov, A., McLean, J.S., Pevzner, P.A.: hybridspades: an algorithm for hybrid assembly of short and long reads. Bioinformatics 32(7), 1009–1015 (2015)

2. Bankevich, A., Nurk, S., Antipov, D., Gurevich, A.A., Dvorkin, M., Kulikov, A.S., Lesin, V.M., Nikolenko, S.I., Pham, S., Prjibelski, A.D., et al.: Spades: a new genome assembly algorithm and its applications to single-cell sequencing. Journal of computational biology 19(5), 455–477 (2012)

3. Chikhi, R., Rizk, G.: Space-efficient and exact de bruijn graph representation based on a bloom filter. Algorithms for Molecular Biology 8(1), 22 (2013)

4. Chin, C.S., Peluso, P., Sedlazeck, F.J., Nattestad, M., Concepcion, G.T., Clum, A., Dunn, C., O’Malley, R., Figueroa-Balderas, R., Morales-Cruz, A., et al.: Phased diploid genome assembly with single-molecule real-time sequencing. Nature methods 13(12), 1050 (2016)

5. Di Genova, A., Buena-Atienza, E., Ossowski, S., Sagot, M.F.: Wengan: Efficient and high quality hybrid de novo assembly of human genomes. bioRxiv p. 840447 (2019)

6. Haghshenas, E., Sahinalp, S.C., Hach, F.: lordfast: sensitive and fast alignment search tool for long noisy read sequencing data. Bioinformatics 35(1), 20–27 (2018)

7. Huang, W., Li, L., Myers, J.R., Marth, G.T.: Art: a next-generation sequencing read simulator. Bioinformatics 28(4), 593–594 (2011)

8. Jaworski, C.C., Allan, C.W., Matzkin, L.M.: Chromosome-level hybrid de novo genome assemblies as an attainable option for non-model organisms. bioRxiv p. 748228 (2019)

9. Jiang, J.B., Quattrini, A.M., Francis, W.R., Ryan, J.F., Rodríguez, E., McFadden, C.S.: A hybrid de novo assembly of the sea pansy (renilla muelleri) genome. GigaScience 8(4), giz026 (2019)

10. Kadobianskyi, M., Schulze, L., Schuelke, M., Judkewitz, B.: Hybrid genome assembly and annotation of danionella translucida. BioRxiv p. 539692 (2019)

11. Kokot, M., Długosz, M., Deorowicz, S.: Kmc 3: counting and manipulating k-mer statistics. Bioinformatics 33(17), 2759–2761 (2017)

12. Kolmogorov, M., Yuan, J., Lin, Y., Pevzner, P.A.: Assembly of long, error-prone reads using repeat graphs. Nature biotechnology 37(5), 540 (2019)

13. Koren, S., Schatz, M.C., Walenz, B.P., Martin, J., Howard, J.T., Ganapathy, G., Wang, Z., Rasko, D.A., McCombie, W.R., Jarvis, E.D., et al.: Hybrid error correction and de novo assembly of single-molecule sequencing reads. Nature biotechnology 30(7), 693 (2012)

14. Koren, S., Walenz, B.P., Berlin, K., Miller, J.R., Bergman, N.H., Phillippy, A.M.: Canu: scalable and accurate long-read assembly via adaptive k-mer weighting and repeat separation. Genome research 27(5), 722–736 (2017)

15. Lee, C.: Generating consensus sequences from partial order multiple sequence alignment graphs. Bioinformatics 19(8), 999–1008 (2003)

16. Lee, C., Grasso, C., Sharlow, M.F.: Multiple sequence alignment using partial order graphs. Bioinformatics 18(3), 452–464 (2002)

17. Li, H.: Minimap and miniasm: fast mapping and de novo assembly for noisy long sequences. Bioinformatics 32(14), 2103–2110 (2016)

18. Li, H.: Minimap2: pairwise alignment for nucleotide sequences. Bioinformatics 34(18), 3094–3100 (2018)

19. Miga, K.H., Koren, S., Rhie, A., Vollger, M.R., Gershman, A., Bzikadze, A., Brooks, S., Howe, E., Porubsky, D., Logsdon, G.A., et al.: Telomere-to-telomere assembly of a complete human x chromosome. BioRxiv p. 735928 (2019)

20. Mikheenko, A., Prjibelski, A., Saveliev, V., Antipov, D., Gurevich, A.: Versatile genome assembly evaluation with quast-lg. Bioinformatics 34(13), i142–i150 (2018)

21. Miller, J.R., Delcher, A.L., Koren, S., Venter, E., Walenz, B.P., Brownley, A., Johnson, J., Li, K., Mobarry, C., Sutton, G.: Aggressive assembly of pyrosequencing reads with mates. Bioinformatics 24(24), 2818–2824 (2008)

22. Molnar, M., Haghshenas, E., Ilie, L.: Sage2: parallel human genome assembly. Bioinformatics 34(4), 678–680 (2017)

23. Myers, G.: Efficient local alignment discovery amongst noisy long reads. In: International Workshop on Algorithms in Bioinformatics. pp. 52–67. Springer (2014)

24. Ono, Y., Asai, K., Hamada, M.: Pbsim: Pacbio reads simulatortoward accurate genome assembly. Bioinformatics 29(1), 119–121 (2012)

25. Ruan, J., Li, H.: Fast and accurate long-read assembly with wtdbg2. BioRxiv p. 530972 (2019)

26. Simão, F.A., Waterhouse, R.M., Ioannidis, P., Kriventseva, E.V., Zdobnov, E.M.: Busco: assessing genome assembly and annotation completeness with single-copy orthologs. Bioinformatics 31(19), 3210–3212 (2015)

27. Simpson, J.T., Durbin, R.: Efficient de novo assembly of large genomes using compressed data structures. Genome research 22(3), 549–556 (2012)

28. Simpson, J.T., Wong, K., Jackman, S.D., Schein, J.E., Jones, S.J., Birol, I.: Abyss: a parallel assembler for short read sequence data. Genome research 19(6), 1117–1123 (2009)

29. Vaser, R., Sović, I., Nagarajan, N., Šikić, M.: Fast and accurate de novo genome assembly from long uncorrected reads. Genome research 27(5), 737–746 (2017)

30. Vollger, M.R., Logsdon, G.A., Audano, P.A., Sulovari, A., Porubsky, D., Peluso, P., Concepcion, G.T., Munson, K.M., Baker, C., Sanders, A.D., et al.: Improved assembly and variant detection of a haploid human genome using single-molecule, high-fidelity long reads. BioRxiv p. 635037 (2019)

31. Walker, B.J., Abeel, T., Shea, T., Priest, M., Abouelliel, A., Sakthikumar, S., Cuomo, C.A., Zeng, Q., Wortman, J., Young, S.K., et al.: Pilon: an integrated tool for comprehensive microbial variant detection and genome assembly improvement. PloS one 9(11), e112963 (2014)

32. Wick, R.R., Judd, L.M., Gorrie, C.L., Holt, K.E.: Unicycler: resolving bacterial genome assemblies from short and long sequencing reads. PLoS computational biology 13(6), e1005595 (2017)

33. Wick, R.R., Schultz, M.B., Zobel, J., Holt, K.E.: Bandage: interactive visualization of de novo genome assemblies. Bioinformatics 31(20), 3350–3352 (2015)

34. Ye, C., Hill, C.M., Wu, S., Ruan, J., Ma, Z.S.: Dbg2olc: efficient assembly of large genomes using long erroneous reads of the third generation sequencing technologies. Scientific reports 6, 31900 (2016)

35. Zerbino, D.R., Birney, E.: Velvet: algorithms for de novo short read assembly using de bruijn graphs. Genome research 18(5), 821–829 (2008)

36. Zimin, A.V., Puiu, D., Luo, M.C., Zhu, T., Koren, S., Marçais, G., Yorke, J.A., Dvořák, J., Salzberg, S.L.: Hybrid assembly of the large and highly repetitive genome of aegilops tauschii, a progenitor of bread wheat, with the masurca mega-reads algorithm. Genome research 27(5), 787–792 (2017)

