## Supplementary material for "HASLR: Fast Hybrid Assembly of Long Reads"

E. Haghshenas et al.

### S1 Data

#### S1.1 Simulated data

PBSIM has an option to infer the mean and standard deviation of read length and the error rate from a real dataset. So first, we prepare that real dataset. We use the first 10 runs of CHM1 (P6C4) dataset:

```
$ for acc in SRR2183739 SRR2183740 SRR2183741 SRR2183742 SRR2183743 SRR2183744 SRR2183745
    SRR2183746 SRR2183747 SRR2183748; do wget http://sra-download.ncbi.nlm.nih.gov/srapub_files/${
    acc}_${acc}_hdf5.tgz; done

$ for acc in SRR2183739 SRR2183740 SRR2183741 SRR2183742 SRR2183743 SRR2183744 SRR2183745
    SRR2183746 SRR2183747 SRR2183748; do tar -zxvf ${acc}_${acc}_hdf5.tgz; done

$ for bax in m15051*.bax.h5; do bash5tools.py ${bax} --outFilePrefix ${bax} --outType fastq --
    readType subreads --minLength 50 --minReadScore 0.75; done

$ for seq in m15051*.fastq; do cat ${seq}; done > chm1_p6c4_first_10.fastq
```

For simulation of the long reads:

```
$ pbsim --seed 0 --data-type CLR --depth 50 --length-min 1 --length-max 500000 --sample-fastq
    chm1_p6c4_first_10.fastq --prefix long <reference_fasta>
```

For simulation of the short reads:

```
$ art_illumina --paired --in <reference_fasta> --len 150 --mflen 500 --sdev 50 --fcov 50 --rndSeed
    0 --noALN --out short
```

#### S1.2 Real data

Table S1: Availability of real long read datasets

| Dataset | Technology | Accession |
|---|---|---|
| *E.coli* | ONT R9.4<br>Illumina | http://lab.loman.net/2017/03/09/ultrareads-for-nanopore<br>ftp://webdata:webdata@ussd-ftp.illumina.com/Data/SequencingRuns/MG1655/MiSeq_Ecoli_MG1655_110721_PF_R1.fastq.gz<br>ftp://webdata:webdata@ussd-ftp.illumina.com/Data/SequencingRuns/MG1655/MiSeq_Ecoli_MG1655_110721_PF_R2.fastq.gz |
| Yeast | PacBio<br>Illumina | ERX1725434, ERX1725435, ERX1725441<br>ERX1943903 |
| *C.elegans* | PacBio<br>Illumina | https://github.com/PacificBiosciences/DevNet/wiki/C.-elegans-data-set<br>SRR065390 |
| Human | PacBio<br>Illumina | https://trace.ncbi.nlm.nih.gov/Traces/sra/?study=SRP044331<br>SRX652547 |

#### S2 Software

Table S2: Version, reference, and respository of utilized software.

| Tool | Version | Reference | Repository |
|---|---|---|---|
| Minia | 3.2.1 | [2] | github.com/GATB/minia |
| minimap2 | 2.17 | [6] | github.com/lh3/minimap2 |
| SPOA | 1.1.3 | [10] | github.com/rvaser/spoa |
| GNU Time | 1.9 | – | ftp.gnu.org/gnu/time/ |
| ART | 2.5.8 | [4] | niehs.nih.gov/research/resources/software/biostatistics/art/ |
| PBSIM | 7fdcefd | [8] | github.com/yukiteruono/pbsim |
| Canu | 1.8 | [5] | github.com/marbl/canu |
| wtdbg2 | 2.5 | [9] | github.com/ruanjue/wtdbg2 |
| hybridSPAdes | 3.13.1 | [1] | github.com/ablab/spades |
| Unicycler | 0.4.8 | [11] | github.com/rrwick/unicycler |
| DBG2OLC | 0246e46 | [13] | github.com/yechengxi/dbg2olc |
| Masurca | 3.3.1 | [14] | github.com/alekseyzimin/masurca |
| Wengan | v0.1 | [3] | github.com/adigenova/wengan |
| QUAST | 5.0.2 | [7] | github.com/ablab/quast |
| BUSCO | 4.0.1 | [26] | busco.ezlab.org |

#### S3 Command details

- Running HASLR

```
$ python3 haslr.py --threads <cores> --type <pacbio|nanopore> --cov-lr 25 --minia-kmer 55 --
    minia-solid 3 --aln-block 500 --out <output_directory> --genome <genome_size> --long <
    lr_file> --short <sr_file_1> <sr_file_2>
```

- Running Canu

```
$ canu -p <assembly_prefix> -d <output_directory> genomeSize=<genome_size> -pacbio-raw <
    lr_file> useGrid=false
```

- Running wtdbg2

```
$ perl wtdbg2.pl -t <cores> -x <rs|ont> -g <genome_size> -o <assembly_prefix> <lr_file>
```

- Running hybridSPAdes

```
$ spades.py -t <cores> -m <max_memory> -1 <sr_file_1> -2 <sr_file_2> --pacbio <lr_file> -o <
    output_directory>
```

- Running Unicycler

```
$ unicycler -t <cores> --no_rotate --no_miniasm --no_pilon -o <assembly_prefix> -1 <
    sr_file_1> -2 <sr_file_2> -l <lr_file>
```

- Running DBG2OLC (based on suggestions on the github repository)

```
$ fastutils interleave -q -1 <sr_file_1> -2 <sr_file_2> | fastutils subsample -q -d 50 -g <
    genome_size> > short.50x.fastq

$ fastutils subsample -l -d 30 -g <genome_size> -i <lr_file> > long.30x.fasta

$ SparseAssembler LD 0 k 51 g 15 NodeCovTh 1 EdgeCovTh 0 GS <genome_size> f short.50x.fastq

$ DBG2OLC k 17 AdaptiveTh 0.01 KmerCovTh 2 MinOverlap 20 RemoveChimera 1 Contigs Contigs.txt
     f long.30x.fasta

$ cat Contigs.txt long.30x.fasta > ctg_pb.fasta

$ ulimit -n 4000

$ split_and_run_sparc.sh backbone_raw.fasta DBG2OLC_Consensus_info.txt ctg_pb.fasta ./
    consensus_dir
```

- Running Masurca
  Content of config.txt

```
DATA
PE= pe <insert_mean> <insert_std> <sr_file_1> <sr_file_1>
PACBIO=<lr_file>
# NANOPORE=<lr_file>
END

PARAMETERS
GRAPH_KMER_SIZE = auto
LHE_COVERAGE=25
CA_PARAMETERS = cgwErrorRate=0.15
KMER_COUNT_THRESHOLD = 1
CLOSE_GAPS=0
NUM_THREADS = <cores>
JF_SIZE = 200000000
END
```

Command

```
bash assemble.sh
```

- Running Wengan

```
perl wengan.pl -t <cores> -a M -p <assembly_prefix> -x <pacraw|ontraw> -g <genome_size> -s <
    sr_file_1>,<sr_file_1> -l <lr_file>
```

#### S4 Identification of unique short read contigs

In order to measure the efficacy of our approach for identifying unique short read contigs, we conducted a set of experiments as follows. First, we simulated a short read dataset based on six different reference genomes: E. coli, Yeast, C. elegans, A. thaliana, D. melanogaster, and GRCh38 human reference genome. For each genome, we used ART [4] to simulate 50× coverage short Illumina reads (2×100 bp long, 500 bp insert size mean, and 50 bp insert size deviation) using the Illumina HiSeq 2000 error model. Next, we used Minia to assemble the simulated short reads using $k$-mer size 49. Finally, to form the ground truth for copy count of each SRC, we mapped the assembled SRCs to the reference genome using Minimap2.

For identification of unique short read contigs, we use the notion of mean $k$-mer frequency of short read contigs as follows. We calculate the mean and standard deviation of $k$-mer frequency of 30 longest contigs ($f_{avg}$, $f_{std}$). At the end, every short read contig whose mean $k$-mer frequency is below $f_{avg} + 3f_{std}$ is considered as unique contig.

Here, we report the precision and recall of the above mentioned approach in identifying unique short read contigs. For each dataset, we evaluate the performance of our approach in identifying unique short read contigs that are longer than a threshold. The length threshold that is used to discard small contigs in this experiment changes from 100 to 1000 with a step size of 100.

As it can be seen in Figure S1, the precision of the identified unique short read contigs is always high regardless of the length threshold. In addition, in all the experiments a big jump in recall is observed at length threshold of 300. Also, with a length threshold higher than 500 we are able to identify the unique contigs with very low false positive rate and higher genome coverage. The results of this experiment shows that proposed approach for identifying unique short read contigs performs well with high precision and recall.

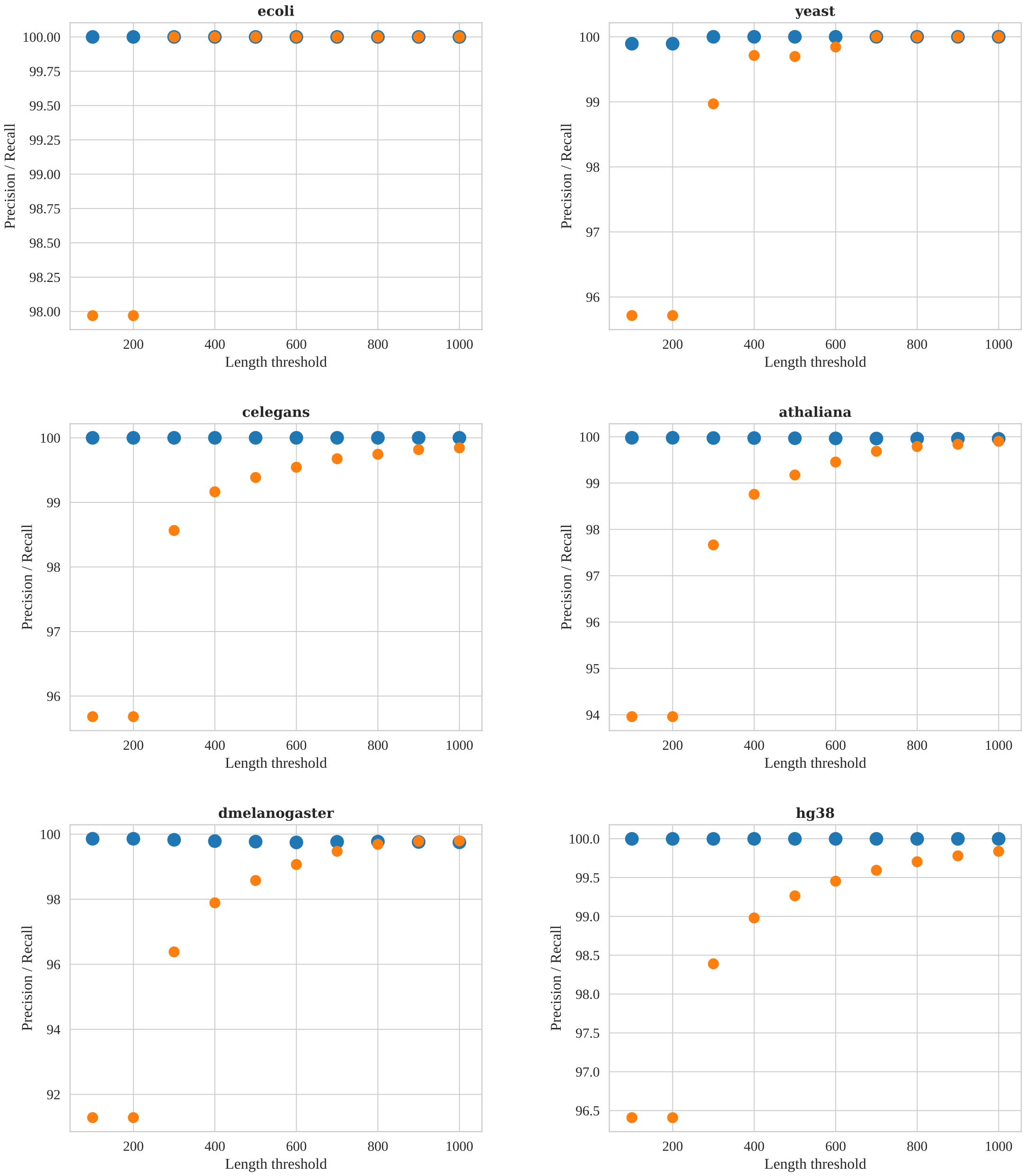


Fig. S1: Precision and recall results in identification of unique short read contigs on 6 different reference genomes. Precision is shown with blue dots and recall is shown with orange dots. Precision is always high across the different experiments and in all the experiments a big jump in recall happens at length threshold of 300.

#### S5 Removing tips and bubbles from backbone graph

In this section we explain the algorithms for removing tips and bubbles from the backbone graph. Tips are dead-end simple paths whose length are small compared to their *parallel* paths. Bubbles are formed when two disjoint simple paths occur between two nodes. Figure S2 shows examples of tips and bubbles in our backbone graph.

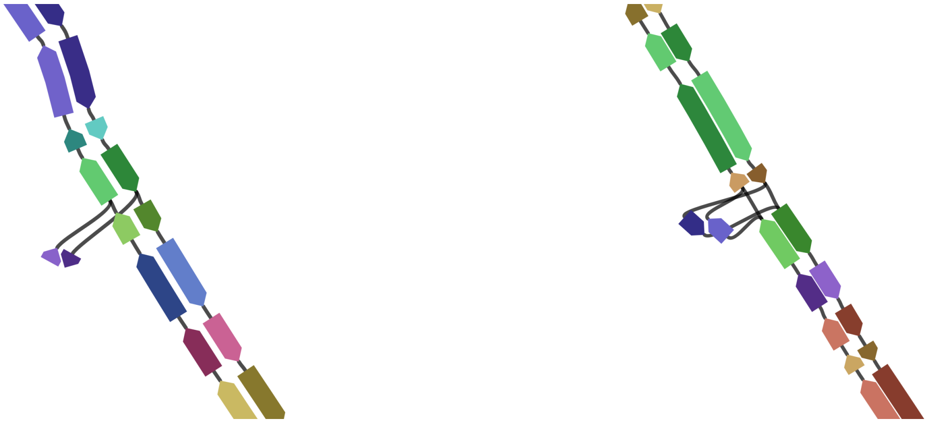

(a) example of a tip in the backbone graph (b) example of a bubble in the backbone graph

Fig. S2: Examples of tip and bubbles in the backbone graph. Here the backbone graph is visualized using Bandage [12]

**Estimation of length and coverage for simple paths.** In order to perform tip and bubble removal, HASLR requires an estimate for the length and coverage of each simple path. Here, we explain how this estimation is calculated.

For each UC in a simple path, we can calculate the coordinates of region that is aligned to all long reads (we refer to this region as *shared* region). Since the length of shared regions corresponding to all UCs are known, we only need to find an estimation for the middle regions (between two consecutive shared regions). To do this, for each long read supporting the edge connecting two UCs, we calculate the length of the LR subsequence that falls between shared regions (using the alignment's CIGAR string). See Figure 2 for a toy example. We use the average of length of all these subsequences as the estimation for the region between shared regions. Finally, the length of the simple path can be estimated as the sum of length of all shared regions plus the estimated length of all middle regions.

In addition, the coverage of each simple path can be calculated based on the number of long reads supporting each edge as well as the estimated length of the middle regions between two consecutive shared regions.

**Bubble removal.** On a haploid genome, our identification of unique short read contigs is accurate, bubbles are caused only by incorrect alignment of UCs in the middle of LRs. In this case, the bubble is usually formed by two simple paths with same length while one of them has a significantly lower coverage.

In contrast, in diploid genomes, it is possible to have natural bubbles corresponding to heterozygous regions of the genome. The main characteristic of such bubbles is having similar coverage on two paths forming the bubble. If the region contains a heterozygous insertion or deletion, the length of two simple paths forming the bubble are different. On the other hand, if the region contains an inversion, two paths have the same length. Therefore, looking at length of the two paths forming the bubble is not a good criteria for identification of artificial bubbles. This means, decision making should be solely based on the coverage of two paths.

**Tip removal.** Tips are mainly caused by incorrect alignment of UCs at the extremities of LRs. As a result, the simple path causing the tip is expected to have a small length. In addition, the coverage of such simple path is usually much lower than other simple paths. In our implementation, a simple path is considered as tip if (i) it is a dead-end (only one end is connected to other nodes) and (ii) contains less than 3 UCs. Based on our observations, most of the tips are dead-end simple paths that contain only a single UC.

#### S6 Polishing of the assemblies obtained from two real datasets

Table S3: Effect of polishing assemblies on the small assembly errors of real datasets

| | | Mismtach rate | | Indel rate | |
|---|---|---|---|---|---|
| Dataset | Assembler | draft | polished | draft | polished |
| Yeast (PacBio) | Canu | 8.85 | 7.56 | 7.99 | 7.99 |
| | wtdbg2 | 10.65 | 7.19 | 27.17 | 2.61 |
| | hybridSPAdes | 44.77 | 9.88 | 3.71 | 3.93 |
| | Unicycler | 15.13 | 6.84 | 4.22 | 2.44 |
| | DBG2OLC | 28.37 | 14.42 | 58.43 | 5.51 |
| | Masurca | 11.83 | 8.49 | 5.85 | 9.69 |
| | Wengan | 11.86 | 7.36 | 34.29 | 2.08 |
| | HASLR | 8.13 | 4.33 | 100.64 | 2.05 |
| *C.elegans* (PacBio) | Canu | 65.28 | 65.88 | 58.82 | 29.71 |
| | wtdbg2 | 26.82 | 25.9 | 79.72 | 27.11 |
| | hybridSPAdes | 108.04 | 27.88 | 15.96 | 45.43 |
| | Unicycler | 58.36 | 36.97 | 45.47 | 32.08 |
| | DBG2OLC | 44.75 | 46.50 | 80.61 | 43.52 |
| | Masurca | 49.20 | 30.9 | 23.50 | 31.97 |
| | Wengan | 35.75 | 21.13 | 121.11 | 22.82 |
| | HASLR | 26.08 | 19.61 | 140.40 | 22.92 |

Note: Here polished genomes are obtained after a single round of polishing using Arrow (`github.com/PacificBiosciences/GenomicConsensus`)

#### S7 Visual examples of regions assembled only by HASLR without any misassembly

We manually inspected the assemblies of simulated datasets (*C. elegans* and human) using QUAST's contig browser tool (Icarus). Here, we have include a few examples of the regions that are properly assembled with HASLR but assemblies of other tools are either fragmented or contain misassemblies.

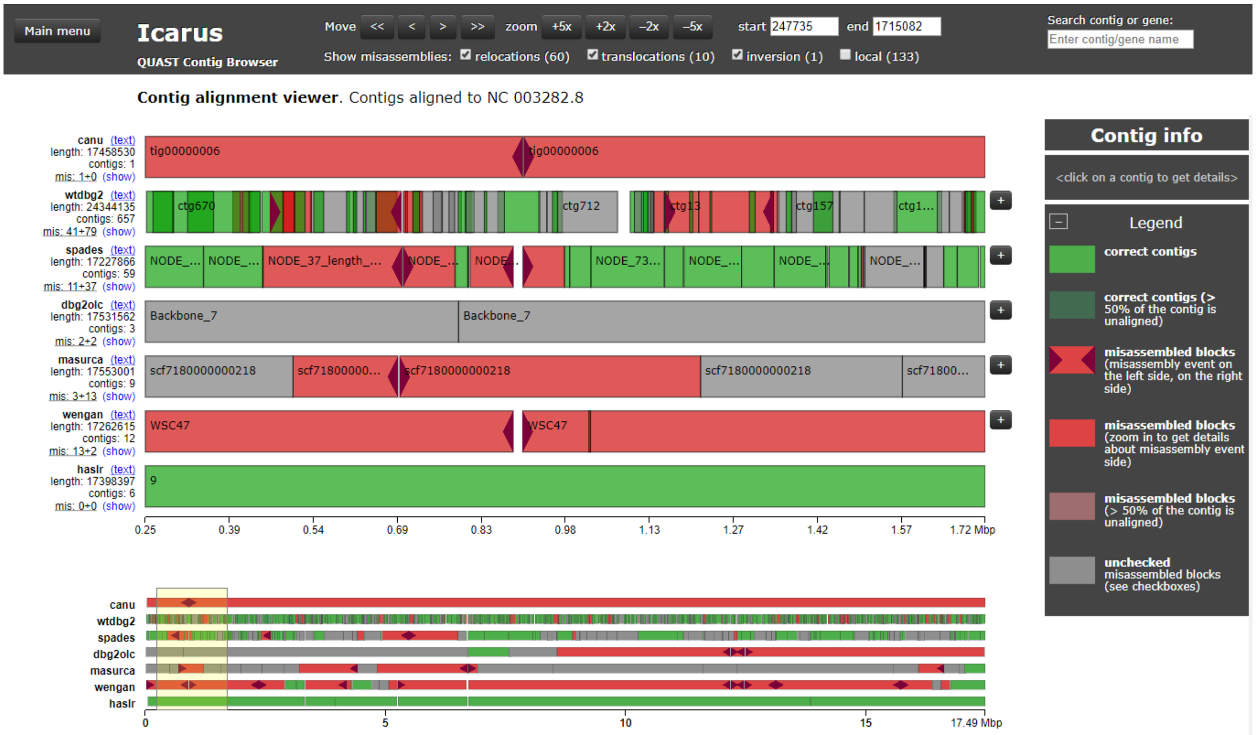


Fig. S3: An example showing a region of choromosome 4 of *C. elegans*.

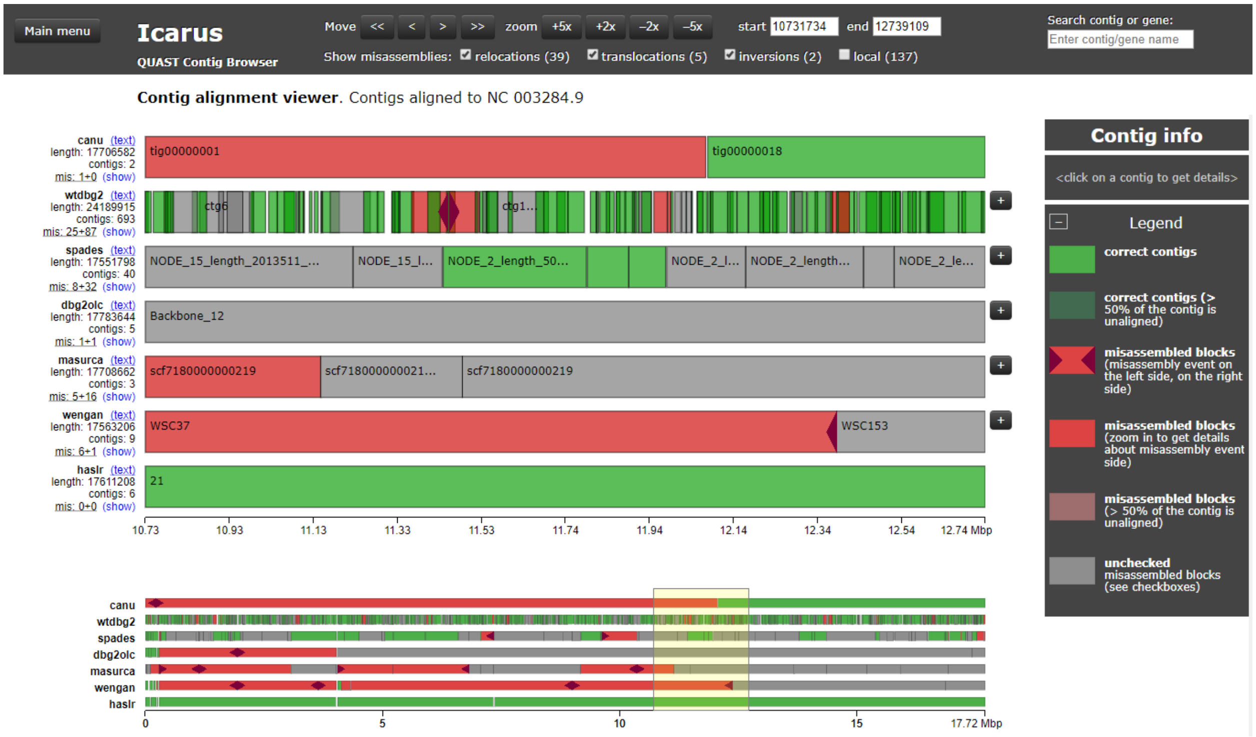


Fig. S4: An example showing a region of choromosome X of *C. elegans*.

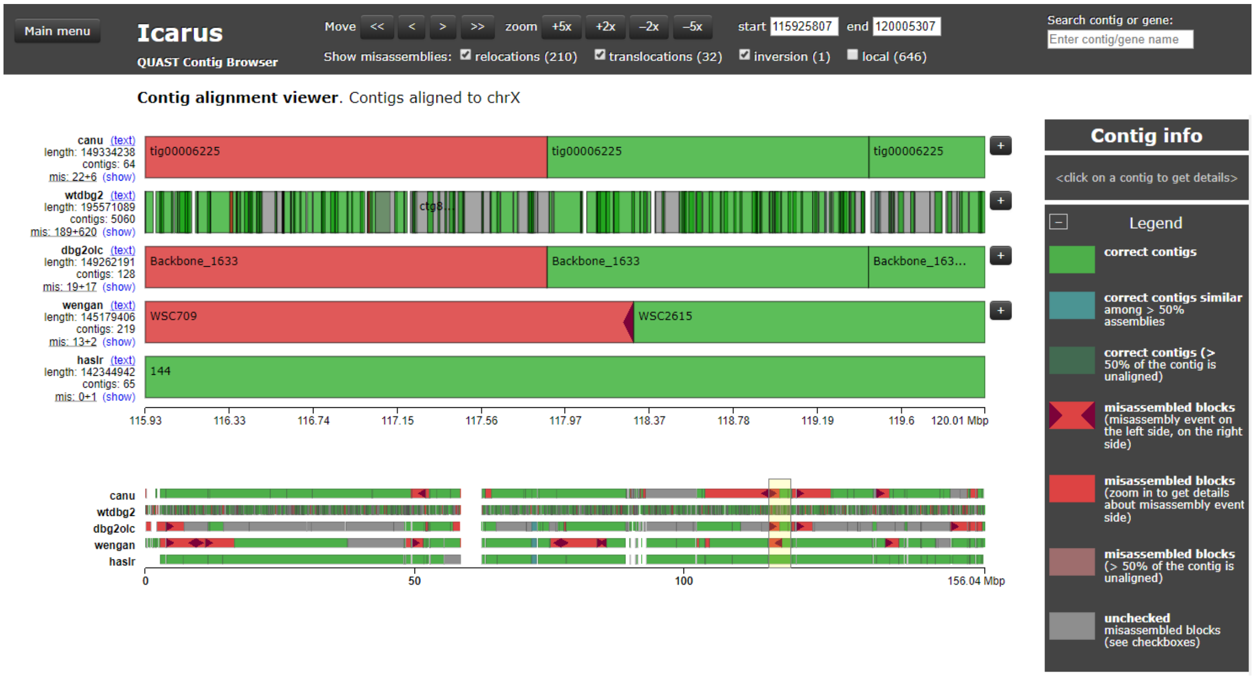


Fig. S5: An example showing a region of choromosome X of hg38.

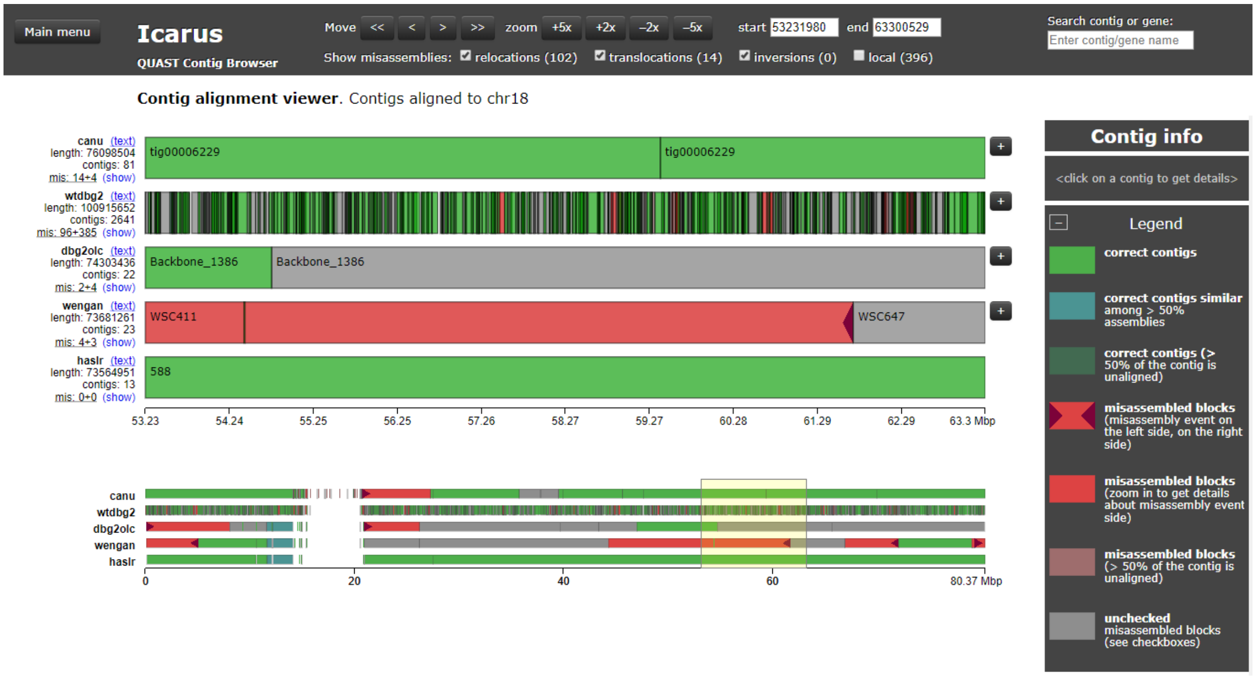


Fig. S6: An example showing a region of choromosome 18 of hg38.

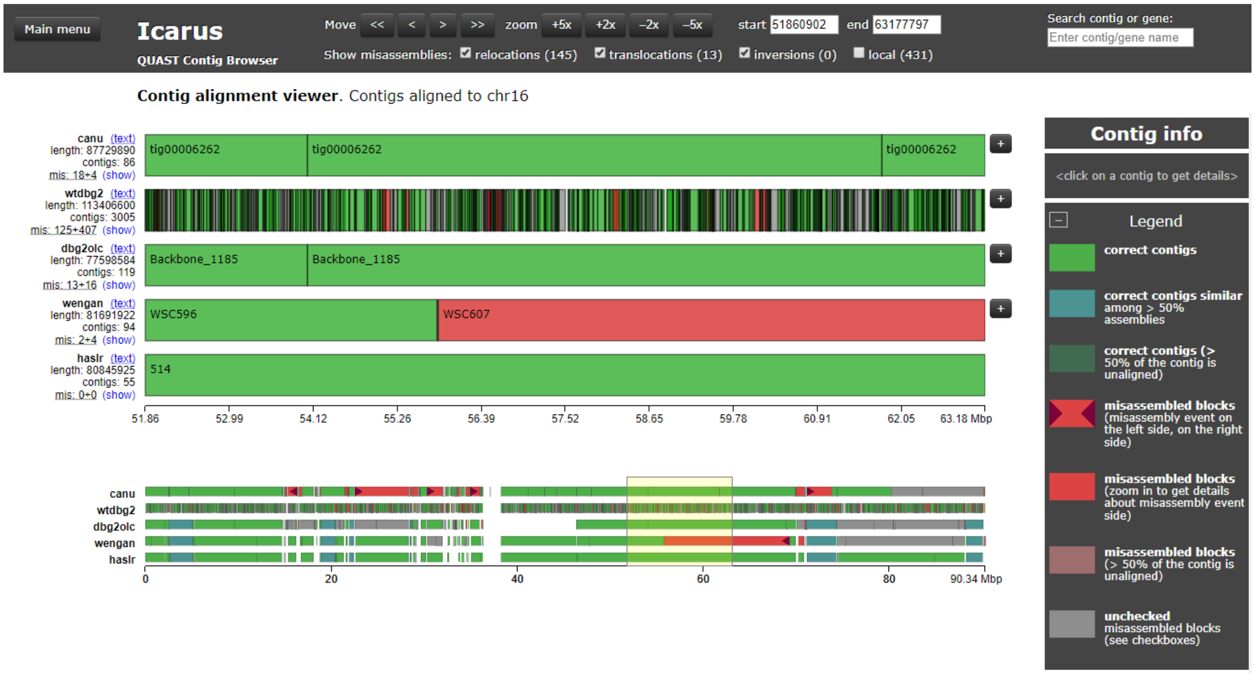


Fig. S7: An example showing a region of choromosome 16 of hg38.

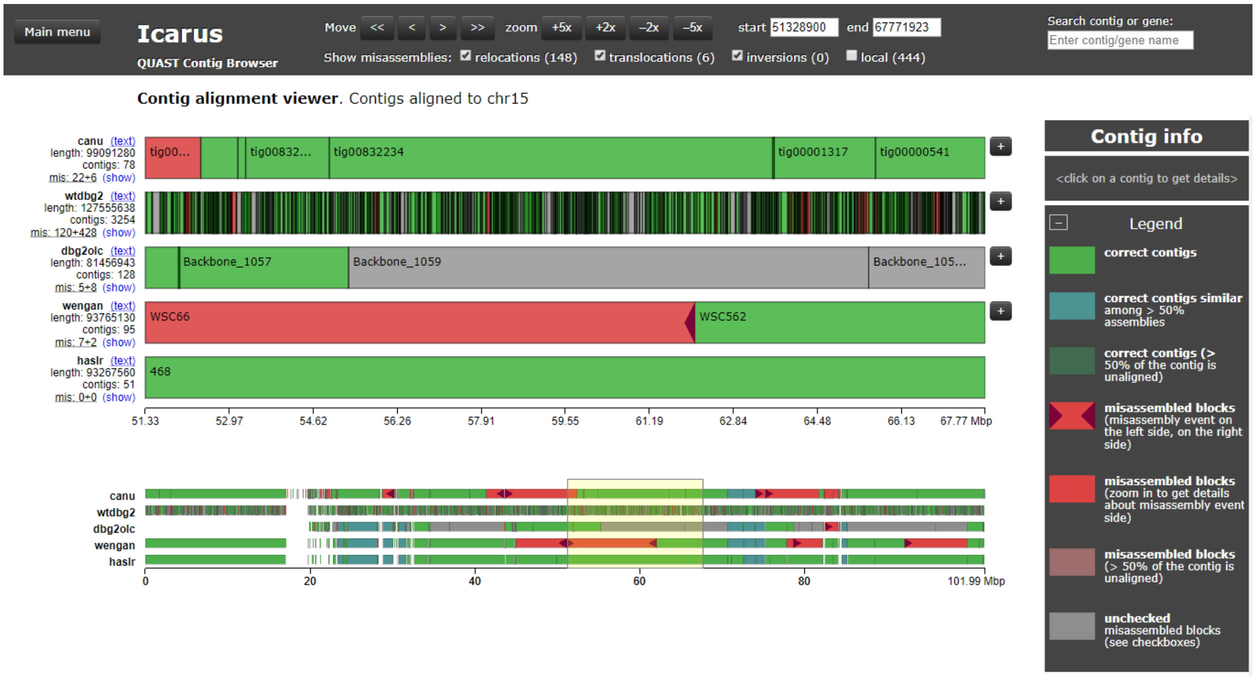


Fig. S8: An example showing a region of choromosome 15 of hg38.

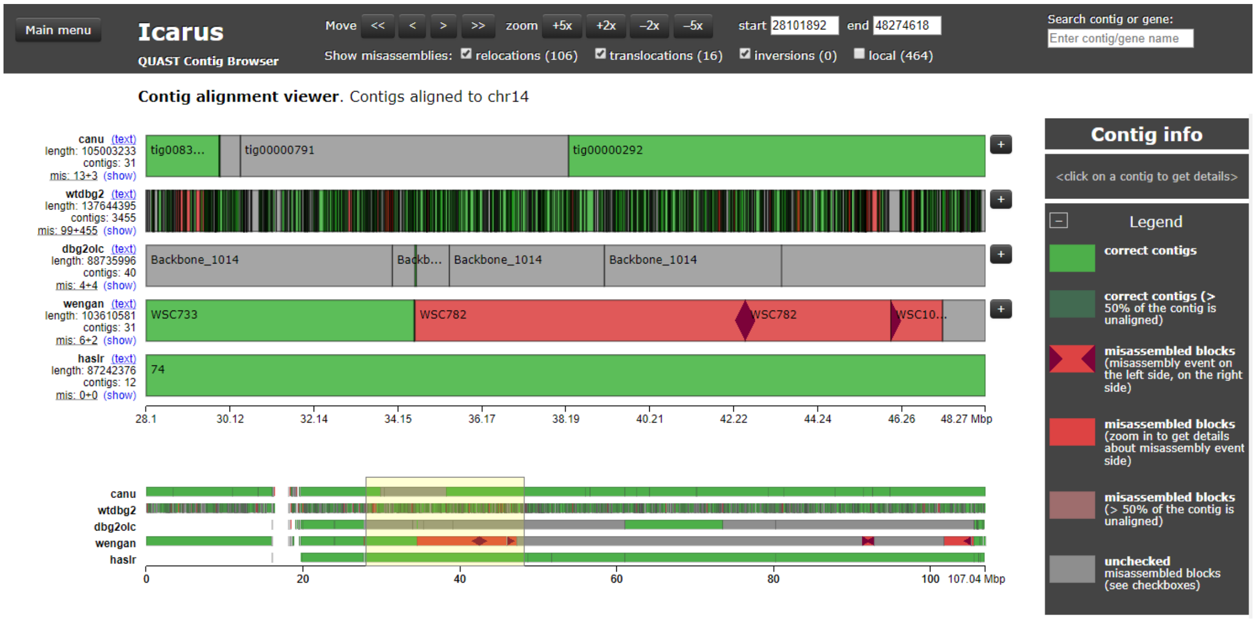


Fig. S9: An example showing a region of choromosome 14 of hg38.

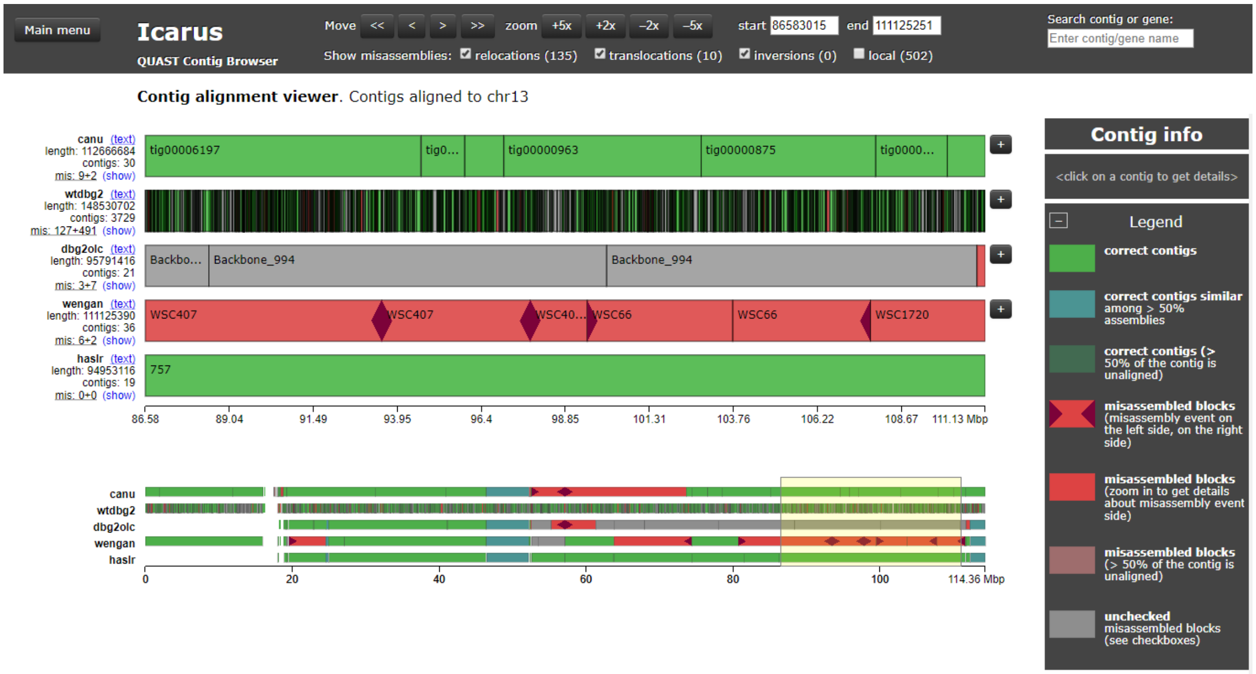


Fig. S10: An example showing a region of choromosome 13 of hg38.

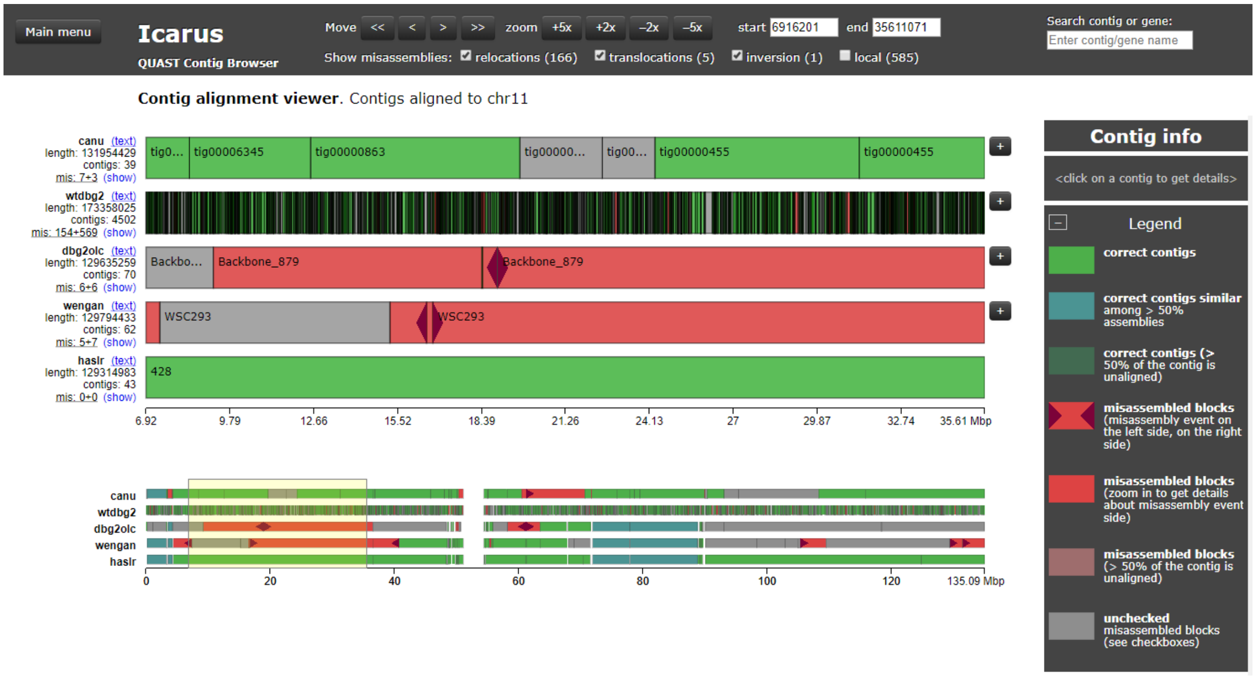


Fig. S11: An example showing a region of choromosome 11 of hg38.

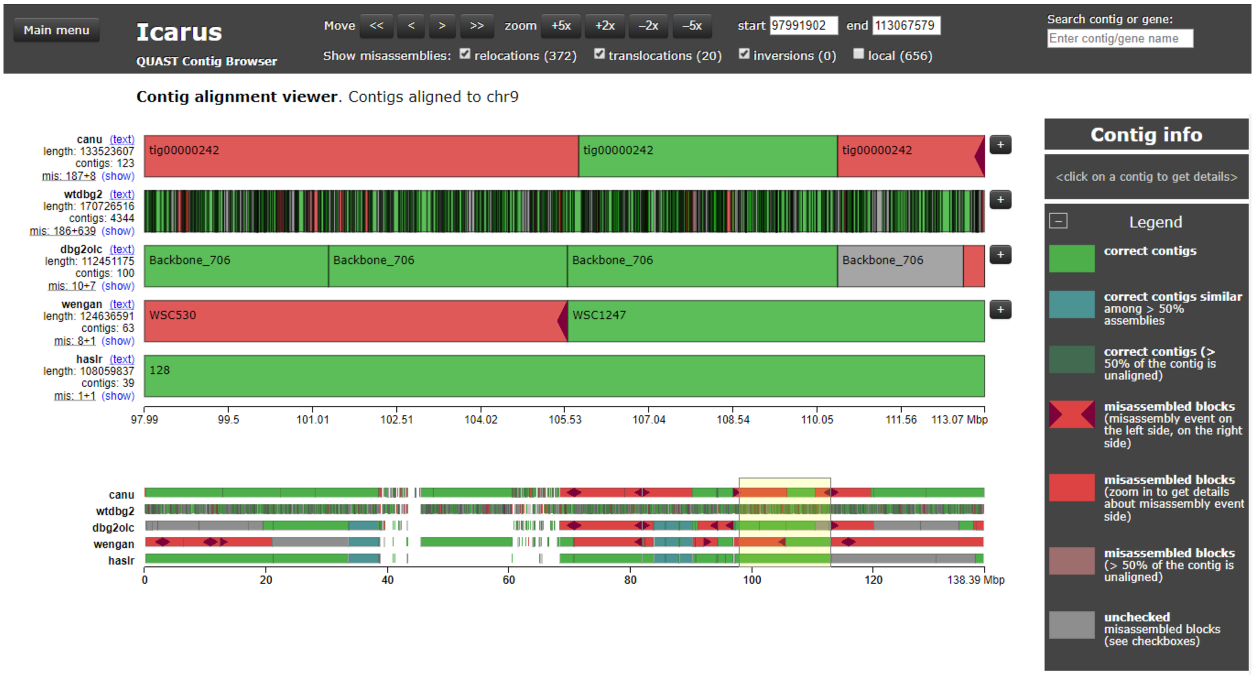


Fig. S12: An example showing a region of choromosome 9 of hg38.

#### References

1. Antipov, D., Korobeynikov, A., McLean, J.S., Pevzner, P.A.: hybridspades: an algorithm for hybrid assembly of short and long reads. Bioinformatics **32**(7), 1009–1015 (2015)
2. Chikhi, R., Rizk, G.: Space-efficient and exact de bruijn graph representation based on a bloom filter. Algorithms for Molecular Biology **8**(1), 22 (2013)
3. Di Genova, A., Buena-Atienza, E., Ossowski, S., Sagot, M.F.: Wengan: Efficient and high quality hybrid de novo assembly of human genomes. bioRxiv p. 840447 (2019)
4. Huang, W., Li, L., Myers, J.R., Marth, G.T.: Art: a next-generation sequencing read simulator. Bioinformatics **28**(4), 593–594 (2011)
5. Koren, S., Walenz, B.P., Berlin, K., Miller, J.R., Bergman, N.H., Phillippy, A.M.: Canu: scalable and accurate long-read assembly via adaptive k-mer weighting and repeat separation. Genome research **27**(5), 722–736 (2017)
6. Li, H.: Minimap2: pairwise alignment for nucleotide sequences. Bioinformatics **34**(18), 3094–3100 (2018)
7. Mikheenko, A., Prjibelski, A., Saveliev, V., Antipov, D., Gurevich, A.: Versatile genome assembly evaluation with quast-lg. Bioinformatics **34**(13), i142–i150 (2018)
8. Ono, Y., Asai, K., Hamada, M.: Pbsim: Pacbio reads simulatortoward accurate genome assembly. Bioinformatics **29**(1), 119–121 (2012)
9. Ruan, J., Li, H.: Fast and accurate long-read assembly with wtdbg2. BioRxiv p. 530972 (2019)
10. Vaser, R., Sović, I., Nagarajan, N., Šikić, M.: Fast and accurate de novo genome assembly from long uncorrected reads. Genome research **27**(5), 737–746 (2017)
11. Wick, R.R., Judd, L.M., Gorrie, C.L., Holt, K.E.: Unicycler: resolving bacterial genome assemblies from short and long sequencing reads. PLoS computational biology **13**(6), e1005595 (2017)
12. Wick, R.R., Schultz, M.B., Zobel, J., Holt, K.E.: Bandage: interactive visualization of de novo genome assemblies. Bioinformatics **31**(20), 3350–3352 (2015)
13. Ye, C., Hill, C.M., Wu, S., Ruan, J., Ma, Z.S.: Dbg2olc: efficient assembly of large genomes using long erroneous reads of the third generation sequencing technologies. Scientific reports **6**, 31900 (2016)
14. Zimin, A.V., Puiu, D., Luo, M.C., Zhu, T., Koren, S., Marçais, G., Yorke, J.A., Dvořák, J., Salzberg, S.L.: Hybrid assembly of the large and highly repetitive genome of aegilops tauschii, a progenitor of bread wheat, with the masurca mega-reads algorithm. Genome research **27**(5), 787–792 (2017)
